# Genotype network intersections promote evolutionary innovation

**DOI:** 10.1101/483040

**Authors:** Devin P. Bendixsen, James Collet, Bjørn Østman, Eric J. Hayden

## Abstract

Evolutionary innovations are qualitatively novel traits that emerge through evolution and increase biodiversity. The genetic mechanisms that lead to innovation remain poorly understood. A systems view of innovation requires the analysis of genotype networks – the vast networks of genetic variants that produce the same phenotype. Innovations can occur at the intersection of two different genotype networks. Here, we use high-throughput sequencing to study the fitness landscape at the intersection of the genotype networks of two catalytic RNA molecules (ribozymes). We determined the ability of numerous neighboring RNA sequences to catalyze two different chemical reactions. We find extensive functional overlap, and over half the genotypes can catalyze both functions to some extent. We demonstrate through evolutionary simulations that these numerous points of intersection facilitate the discovery of a new function. Our study reveals the properties of a fitness landscape where genotype networks intersect, and the consequences for evolutionary innovations.

## MAIN TEXT

### Introduction

The mechanisms by which evolution produces new functions has intrigued biologists since the earliest formulations of evolutionary theory (*1, 2*). Random genetic changes and natural selection would seem to prevent novelty by keeping populations near genotypes at the peaks of fitness landscapes, preserving existing forms at the expense of novel mutants (*3–6*). Models to explain the origins of new functions often invoke gene duplication events, which create redundancy needed to allow either copy to eventually evolve toward a new function (*7–10*). However, the fitness landscape between old and new functions has been difficult to study largely because of the vast number of possible genetic variants for any given gene. As a result, models of innovation differ in the relative importance of neutral drift, environmental changes, the timing and type of selection pressure, and the high-dimensional nature of sequence space (*11*). Our understanding of innovations will benefit from direct observations of the evolution of new functions (*12–17*).

Macromolecular phenotypes such as enzymes can tolerate changes to their primary sequence without necessarily changing structure or function. As a consequence of this robustness to mutations, many genotypes have the same phenotype (*18, 19*). Natural populations of both organisms and macromolecules that appear the same phenotypically still harbor many genetic differences. Genotype networks are the collection of all genotypes with the same phenotype that are interconnected by mutational steps (*20*). Populations occupy finite regions of these vast networks, and it has been suggested that innovations can occur when populations encounter regions of genotype space where two different genotype networks are in close proximity (*21*) (Fig. 1A). To evaluate various models of molecular innovation, it is necessary to characterize the number of mutations that separate two networks and the fitness consequences of the mutational changes needed to move from one network to the other.

**Fig. 1.**
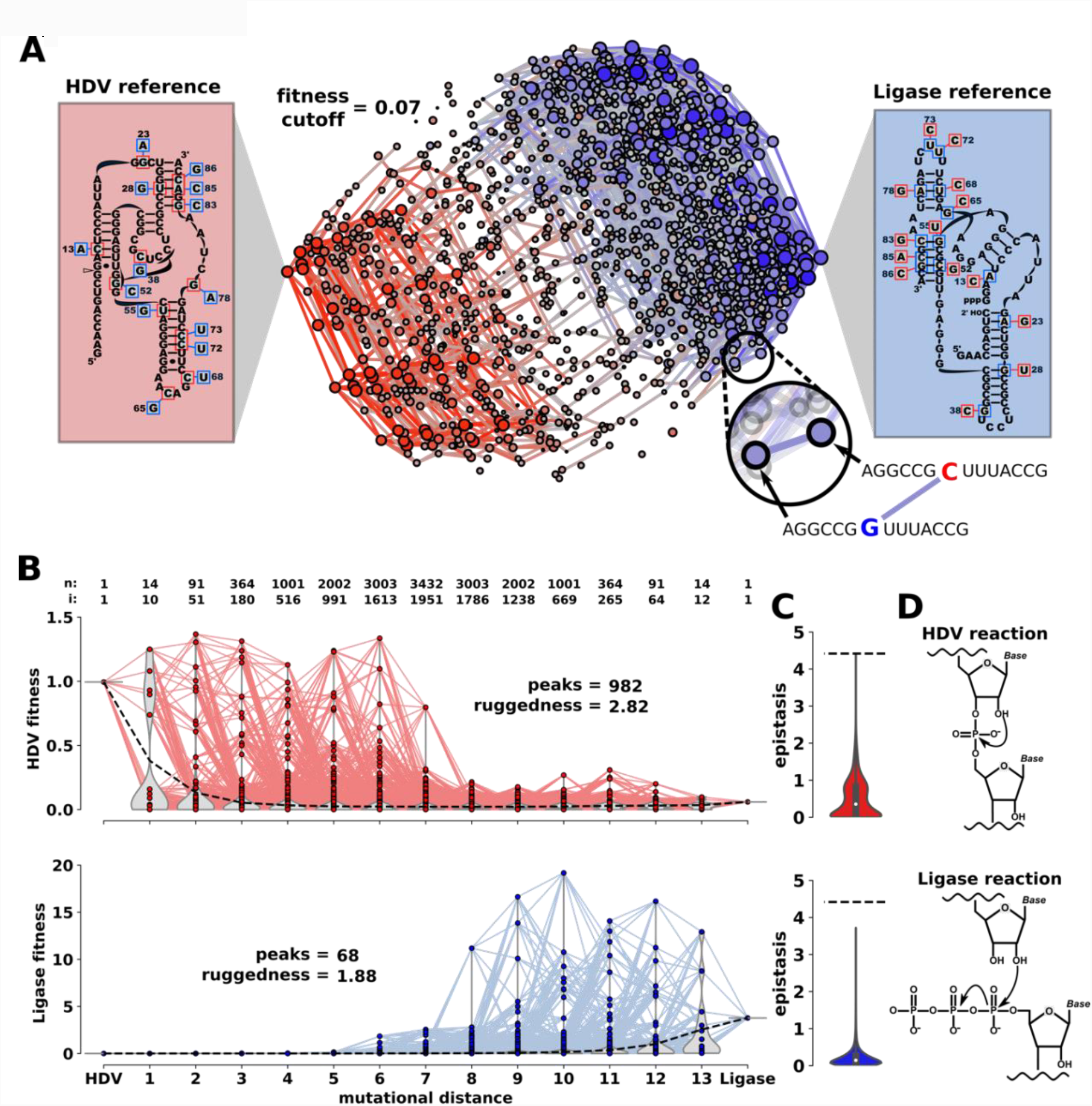
The experimental fitness landscape at the intersection of two genotype networks. **(A)** Overlay of the HDV and Ligase genotype networks. Nodes represent individual genotypes that are connected by an edge if they are different by a single nucleotide change. Nodes are colored based on their dominant activity (red = HDV; blue = Ligase). For each genotype, *fitness* is defined as the relative ribozyme activity determined by high-throughput sequencing and is indicated by the size of the node and the color saturation. Genotypes with fitness below 0.07 are excluded for visualization purposes. Boxes on the left (HDV reference) and right (Ligase reference) show the secondary structure for the reference genotypes, and all the mutational changes that were analyzed. The mutations in blue boxes convert the HDV reference to the Ligase reference. The mutations in red boxes convert the Ligase reference to the HDV reference. **(B)** Distance-based layout of the two fitness landscapes. Each sequence is positioned on the x-axis according to its mutational distance from the HDV reference genotype. HDV fitness (red) and Ligase fitness (blue) are indicated by the y-axis value. The number of genotypes (n) increases in the middle of the plot, and the total number of genotypes at each position is indicated about the graph. The number of dual-function intersection sequences (i) at each mutational distance is also indicated. Inset text *peaks*, and *ruggedness* describe quantitative characteristics of the landscapes. **(C)** Distributions of epistatic values found in each landscape. **(D)** Representation of the chemical reactions catalyzed by each ribozyme.

Here, we report an experimentally constructed intersection of two genotype networks. For our study system we have chosen two distinct RNA phenotypes. The RNA molecules are ribozymes, structured RNA molecules that catalyze chemical reactions. One ribozyme phenotype is the naturally-occurring self-cleaving HDV ribozyme. The second phenotype is the class III Ligase ribozyme that was discovered through artificial selection in a lab (*22*). The two ribozymes share no evolutionary history, fold into very different structures (Fig. 1A) and catalyze different chemical reactions (Fig. 1D). Despite the differences between the two ribozymes, it was previously shown that the two genotype networks come in close proximity, and very few mutations could convert one ribozyme into the other (*23*). This provides an experimentally tractable example of a molecular innovation. To characterize the fitness landscape between the two genotype networks we developed two high-throughput sequencing based assays to quantify both ribozyme phenotypes. We identified two reference genotypes with approximately “wild-type” levels of activity that contained 14 mutational differences between them. We synthesized DNA templates need to transcribe the RNA molecules that contain all the pairwise combinations of these mutational differences. We analyzed the 2^14^= 16,384 neighboring RNA sequence variants using both ribozyme assays. For each sequence, we determined the *ribozyme fitness* for both activities, defined as the catalytic activity relative to a reference sequence. For the HDV phenotype, our fitness is defined as how well each sequence self-cleaves during transcription, and for the Ligase phenotype, fitness is defined as the change in abundance of each sequence from a single round of selection for Ligase activity (see Materials and Methods and SI Appendix). With these fitness values, we analyzed the billions of mutational trajectories between the two genotype networks and used computational simulations to explore how these genotype networks facilitate or inhibit evolutionary innovations.

## Results

### Empirical fitness landscape at the intersection of two genotype networks

We obtained fitness measurements for all 16,384 RNA sequences for both RNA phenotypes. For visualization of the resulting genotype networks, we plot the data as a network graph, where each node is a unique sequence, nodes are connected if they differ by a single mutation, and the fitness is represented by the size of the node (Fig. 1). Each node is colored based on the dominant activity, with HDV in red and Ligase in blue. Fitness values were normalized such that *fitness* = 1 for the reference ribozyme, previously referred to as the “prototype” (*23*). This representation of the data allows a visual appraisal of the proximity of the two genotype networks. In general, both networks are characterized by a decrease in fitness with distance from the reference. The region where the two networks are in closest proximity contains sequences with low activity for either function. Still, we find that numerous genotypes in the two networks are proximal, and numerous distance measurements are required to characterize the mutational distance between the networks.

### Proximity and functional overlap of the two genotype networks

To quantify the average distance between the two genotype networks, we measured the mutational distance between every genotype on one network and the nearest genotype on the other network (Fig. 2A). We find that this distance depends upon whether or not a lower bound is set for genotypes to be considered a member of the genotype network. We find that the average distance between the networks decreases as the fitness cut-off is lowered (Fig. 2A). For example, if “wild-type” activity is required (*fitness* > 1), the two networks are separated by ∼7 mutations on average (fig. S1). However, if molecules with 10% of wild-type activity or better are considered part of the network, then most genotypes are only 1-2 mutations from the other network.

Surprisingly, if we do not set any fitness cut-off, and count all genotypes as being a part of a network as long as they were detected as catalytically active in all three replicates of our assay, we find that over half the molecules can actually perform both functions (Fig 2B). The precise count of intersection sequences depends on our minimum read depth requirement. If we require that each sequence must be detected as active (cleaved/ligated) three times and at least once in each replicate, we find 9,032 intersection sequences. If we require that each sequence needs more than one observation in at least one replicate, we find 8,805 intersection sequences. These requirements were based on the probability of sequence errors in our data (fig. S9C). Most of these dual-function intersection sequences have very low fitness for both functions, and not surprisingly, no single sequence had higher than wild-type fitness for both functions (log10(*fitness*) > 0). However, several sequences do show detectable levels of activity for one function and higher than wild-type fitness for the other function. Under many evolutionary scenarios, these genotypes would be the most likely to facilitate a molecular innovation because they would be favored if selection was acting on only one function, yet would already provide the new function as a suboptimal promiscuous function (*24, 25*). These results demonstrate that the genotype networks have substantial overlap with numerous intersection sequences.

**Fig. 2.**
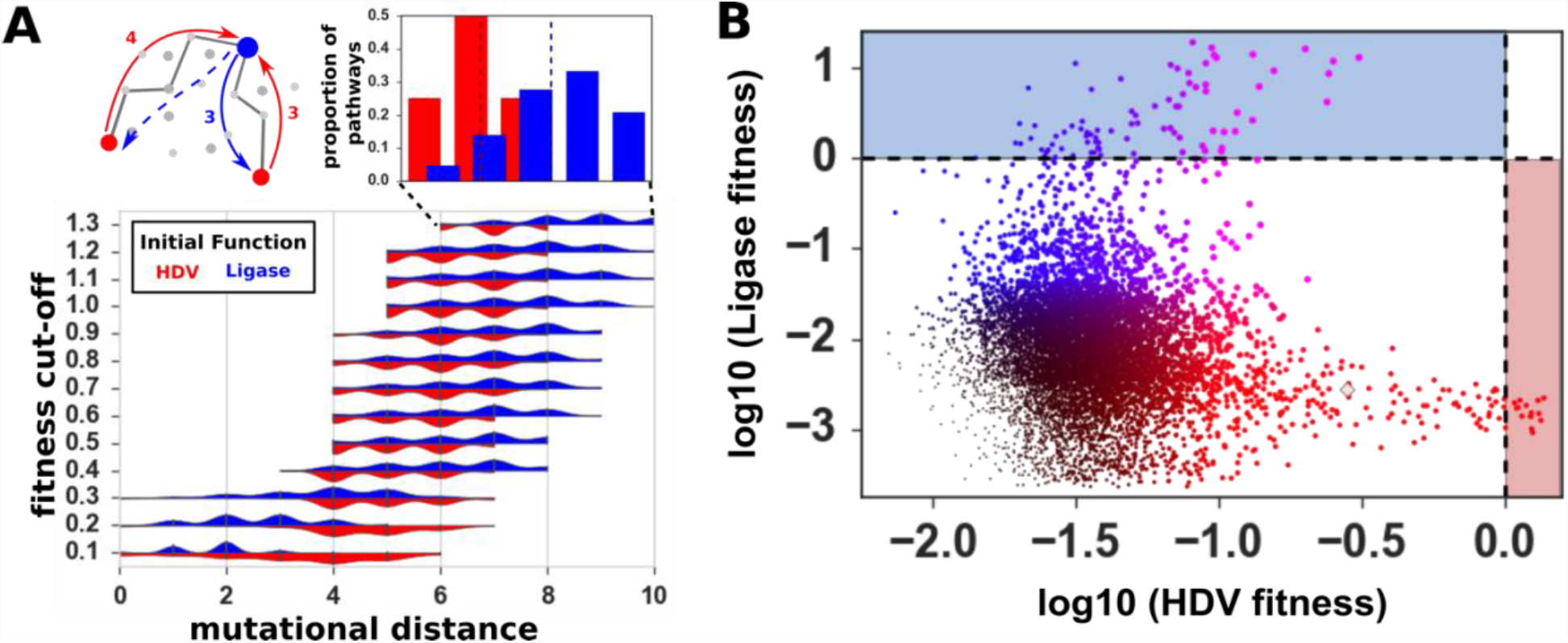
Proximity and overlap of the two genotype networks. **(A)** Distributions of shortest *mutational distance* (x-axis) between genotypes on different networks as a function of *fitness cut-off* (y-axis; blue = Ligase to HDV distances; red = HDV to Ligase distances). For each genotype with a fitness above the cut-off value for one function, the distance to the nearest genotype with the other function was determined. The distribution of these distances determined for all genotypes are plotted as violin plots. The diagram (above, left) illustrates the measurement of distance between the two functions. Inset (above, right) shows the distribution at *fitness cut-off* = 1.3 as histograms, and dashed lines indicate the sample means. **(B)** Intersection sequences with detectable activity for both functions. For each genotype, the HDV fitness is plotted on the x-axis and Ligase fitness is plotted on the y-axis. Color indicates the ratio of Ligation fitness (blue) to HDV fitness (red). The size of the node is scaled to the higher of the two fitness values. Fitness values are log10 transformed. Dashed lines indicate *wild-type* level activity with *fitness = 1* (log10 *fitness*= 0).

### Computational simulations of evolutionary innovation on the empirical fitness landscape

Next, we set out to evaluate the implications of these genotype networks for the evolution of molecular innovations. The networks are in fact high-dimensional, which limits any intuitive interpretation. We therefore turned to computational simulations of populations of RNA molecules evolving on the networks. We modeled evolution using a Wright-Fisher model (*26*) with a fixed population size, a fixed mutation rate, and selection determined by the relative fitness of genotypes (see Materials and Methods). To simulate evolutionary innovations, we imagined the naturally occurring HDV genotype as the established function and the *in vitro* selected Ligase activity as the “new” function. We modeled a situation where the enzymatic function of the HDV ribozyme is first under selection, but gene duplication allows a copy of the gene to evolve under selection for Ligase activity. We therefore apply immediate selection pressure using the Ligase fitness measurements, with no further consequence for the changes in HDV activity. For these simulations, it is useful to visualize the genotype networks as a landscape where the height of the landscape is determined by the fitness (Fig. 3A and movie S1). In our simulations, evolving populations will tend to move uphill towards *peaks*, defined as sequences where all 1-mutation neighbors have lower fitness. The crossing of fitness valleys to get from low-fitness peaks to higher-fitness peaks is allowed in our simulations but requires a stochastic series of less likely events. We started multiple simulations from different genotypes on the HDV network and challenged the populations to evolve on the Ligase fitness landscape. We recorded these simulations as movies to observe the process of evolution toward the *new* Ligase function (Fig. 3B and movie S2-S5).

**Fig. 3.**
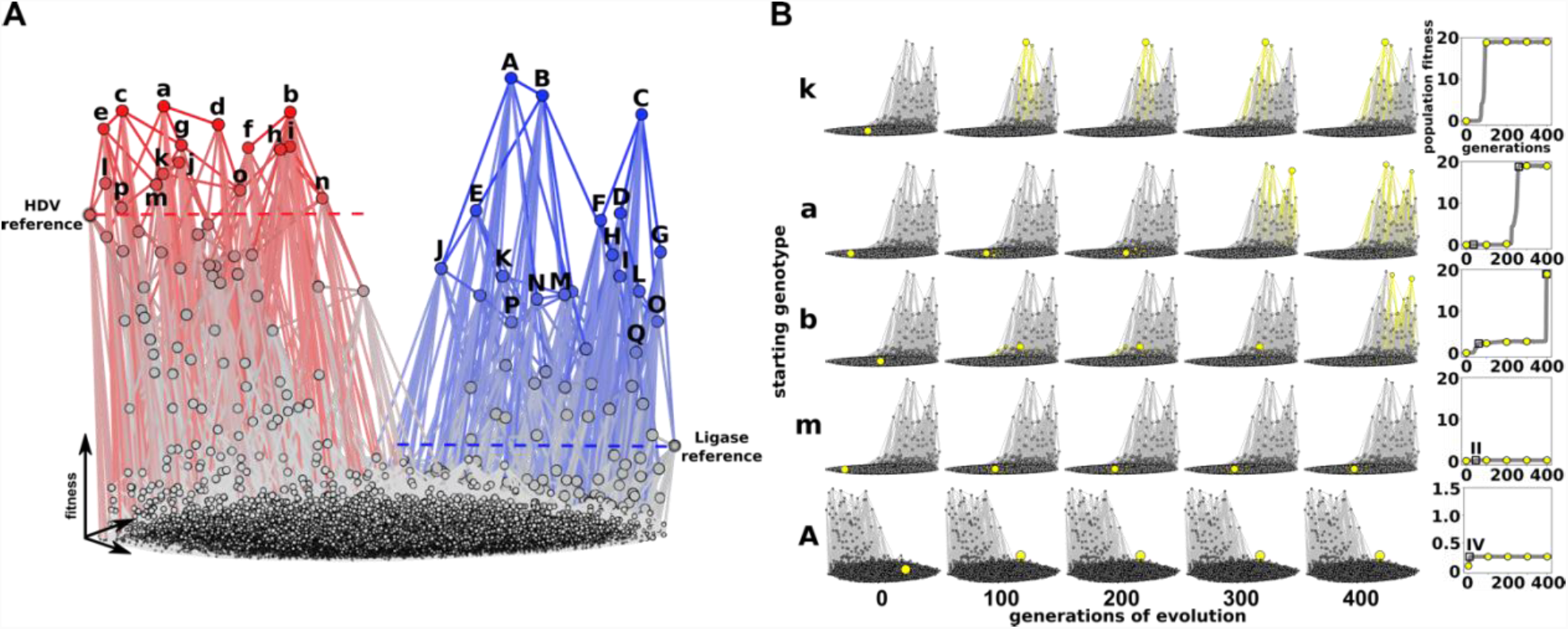
Periods of evolutionary stasis revealed by computational simulation of evolutionary innovation. **(A)** A landscape visualization of the two genotype networks. The height of each node (z-axis) indicates the relative fitness for the HDV phenotype (red) and the Ligase phenotype (blue). Nodes represent genotypes and are connected by an edge if they are different at one nucleotide position. Fitness is indicated by the height (z-axis), the size of the node, and the color saturation. Fitness values are normalized so that both graphs are similar heights. Genotypes used to start evolutionary simulations are labelled with lower case for the genotypes with the highest HDV fitness (a-p) and capital letters for genotypes with the highest Ligase fitness (A-Q). **(B)** Frames from simulations of evolving populations. Several examples are shown to illustrate different rates of increase of *population fitness* over simulation time (*generations*). Each row shows the progress of a single simulation. The starting genotype is indicated to the left. Each plot shows the genotypes present in the population with the number of *generations of evolution* labeled at the bottom. Genotypes present in the population are indicated by yellow nodes and edges. The corresponding *mean fitness* of each population over time is shown in the plots to the right. During simulations the population size (*N*= 1000) and mutation rate (*μ*= 0.01) were constant.

We noticed that many of the individual simulations had periods where the mean fitness of the population plateaus at a specific, often low value for many generations (Fig. 3B). To evaluate the average contribution of these periods of stasis, we repeated the simulation 100 times and plotted the average fitness of the evolving population over time (Fig. 4A, fig. S2). We carried out 100 replicates each for all of the different starting genotypes (Fig. 4B and table S1). We found that different genotypes on the HDV network resulted in consistently different average rates of adaptation to the *new* Ligase function (Fig. 4D). The fact that some genotypes promoted very rapid adaptation supports the idea that *neutral* evolution that enables a population to explore a genotype network can facilitate evolutionary innovations (*21, 27*).

Additionally, we found that there exist specific genotypes on the Ligase fitness landscape that caused these periods of stasis and slower average rates of adaptation (Fig. 4F, fig. S3–S4). These genotypes are local peaks that are characterized by very few pathways to higher fitness. Importantly, the genotypes that caused the slowest rates of adaptation are characterized by extensive reciprocal sign epistasis, meaning that achieving higher fitness requires two or more mutational steps, but *every* initial step is deleterious. Specific starting genotypes on the HDV network frequently stalled at the same intermediate fitness level indicating that they were likely to encounter a specific stasis-causing fitness peak. The consistent rates of adaptation from multiple simulations are encouraging for efforts aimed at forecasting evolutionary outcomes, especially in cases where the underlying fitness landscape can be measured or accurately estimated (*28, 29*).

We next repeated the evolutionary simulations from the opposite perspective, starting with genotypes from the Ligase side of the landscape with selection for improved HDV function. This scenario models Ligase as the original function, and HDV self-cleavage as the new function that is under selection following gene duplication. Surprisingly, we found that all of these simulations got stuck at very low fitness for the full 1000 generations (Fig. 4C, fig. S5), resulting in significant slower rates of adaptation (Fig. 4E). We note that the simulations were done under identical population size and mutation rate, and we therefore attribute the different evolutionary dynamics to properties of the fitness landscapes. The property identified that was likely to dictate evolutionary dynamics was the ruggedness of the landscape. We find that the HDV landscape is much more rugged than the Ligase landscape, with more peaks and more extensive sign epistasis. The HDV landscape has 982 peaks while the Ligase landscape has only 68, caused by more frequent instances of sign epistasis. The severity of sign epistasis is also higher on the HDV landscape, which can be seen in the extreme values in Fig. 1C.

### Co-selection model of evolutionary adaptation

We also modelled a scenario where both functions were simultaneously under selection and each function contributed to fitness (fig. S6–S7). For these simulations, we assigned each genotype a fitness that was calculated by summing the HDV and Ligase fitness values, each multiplied by a weighting parameter that could be adjusted. Before summing, we normalized the data by dividing each fitness values by the maximum fitness value in that landscape, such that the maximum fitness of both functions was *fitness* = 1. We found that under this scenario, the function that was weighted more heavily would be optimized at the expense of the function with lower weighting. Interestingly, when both functions were given equal weight, the “choice” was stochastic and either HDV or Ligase could be optimized. We did not observe any instances where the population remained split with some genotypes being selected for high HDV fitness and others with high Ligase fitness. This suggests that prior to gene duplication a given population is likely to have genes that are optimized for one function, or the other, but not both. This scenario assumes a constant environment. Alternatively, a fluctuating environment could alter selection pressures and help maintain both functions (*30, 31*), and a sudden environmental shift could quickly favor one function over the other.

## Discussion

The difference in the evolutionary dynamics on each landscape have implications for evolutionary innovations. For example, our results indicate that the order in which new functions arise can alter evolutionary dynamics, because optimizing HDV activity from sequences with high Ligase activity is more challenging than evolving in the reverse order. In addition, we found that the specific properties of genotype networks can dictate the rate of evolutionary adaptation of the new function. The Ligase landscape is less rugged with fewer peaks which allows the more rapid evolutionary adaptation of the new function. Taken together, we propose that ancient genes that duplicated and enabled radiation events (*32*) may be characterized by both significant functional overlaps and a robust genotype network. This provides insight into how intersection sequences promote the evolution of new functions and enable the expansion of biodiversity. Further investigations into intersection sequences and fitness landscapes will be required to fully evaluate this scenario. For example, our current library design only investigates two nucleotides at each variable position, which represent the parsimonious or “direct” pathways between the two reference genotypes. However, experimental evidence from a protein enzyme supports the idea that higher-dimensional “indirect” pathways can bypass epistasis and facilitate adaptation (*33*). Further experiments are required to determine how higher-dimensional landscapes contribute to evolutionary innovations (*34*).

It is interesting to note that our results are consistent with observations that have been made from the computational folding and evolution of RNA secondary structures. For example, one prior computational study of simple RNA secondary structures, termed “shapes”, looked at the most probable new shapes that are one mutation away from sequences that form a canonical tRNA structure (*35*). The authors found that most single mutations produce very similar shapes. However, they also found that there exists a high-probability that a single mutation will produce shapes with considerable differences. The HDV and Ligase structures in the current study do not share any structural similarity, but our results show that the shapes overlap extensively in sequence space such that there is a high probability of finding one ribozyme in the neighborhood of the other. These results provide a powerful validation of the insight gained from the computational studies. This similarity is somewhat surprising because the ribozyme phenotypes studied here require a precise tertiary structure to achieve catalysis that is not approximated in secondary structure alone. Nevertheless, the canonical base pairing interactions that are predicted computationally make up a large component of the structural interactions needed for ribozyme folding, which may account for the similarity between the results. Regardless, our results further validate the long-standing use of computational prediction of RNA structures as a realistic model of the genotype to phenotype relationship, which continue to inspire experiments. This also provides motivation for continued efforts to use experimental secondary structure probing methods to improve the blind prediction of RNA tertiary structure (*36*).

The decrease in the fitness of both functions at the intersection also suggests that intermediate forms are disfavored over the sequences that can do one function well (*11*). The evolution of innovation in this sequence space is therefore not only possible, but probable because once a population discovers this region of sequence space, selection is likely to favor a genotype on one side or the other. However, it remains unknown if these characteristics are common or peculiar to these specific phenotypes. Further research advancements will be required to study larger expanses of genotype space needed to cover more mutational positions and higher-dimensionality caused by including all four nucleotides at variable positions. It will also be important to investigate if historic evolutionary innovations found in natural systems have properties like the model system studied here. The high probability of finding dual-function sequences in our current data encourages the search for more genotype network intersections and motivates future research on the forecasting of evolutionary innovations.

## Materials and Methods

### Library Design

For our experiments, we first identified an HDV and a Ligase reference sequence (Fig. 1A). For this purpose, we chose sequence variants that were expected to have near wild-type ribozyme fitness and that were 14 mutations apart (*37*). We then set out to construct a library of ribozyme sequences that contained all the possible presence-absence combinations of these 14 nucleotide differences. These sequence variants represent all the parsimonious intermediates on the evolutionary trajectories between the two reference sequences. Library construction was accomplished by chemically synthesizing a degenerate DNA oligonucleotide that would serve as a template for in vitro transcription with T7 RNA polymerase. At each position where the Ligase and HDV reference ribozymes differed, the synthesis used equal mixtures of two nucleotide phosphoramidites, generating approximately equal probability of both sequence variants. This creates 2_14_= 16,384 ribozyme variants. We synthesized two such libraries, one “HDV-library” with a 5’-leader sequence that is cleaved by variants with the HDV phenotype, and a second “Ligase-library” that begins at the 5’-end of the Ligase ribozyme, so that variants with the Ligase phenotype could react with a separate substrate oligonucleotide (*22*). A common sequence was added to the 3’-end of both libraries to serve as a universal primer binding site for reverse transcription (*38*). Oligonucleotides used in this experiment are listed in Table 1.

**Table 1.**
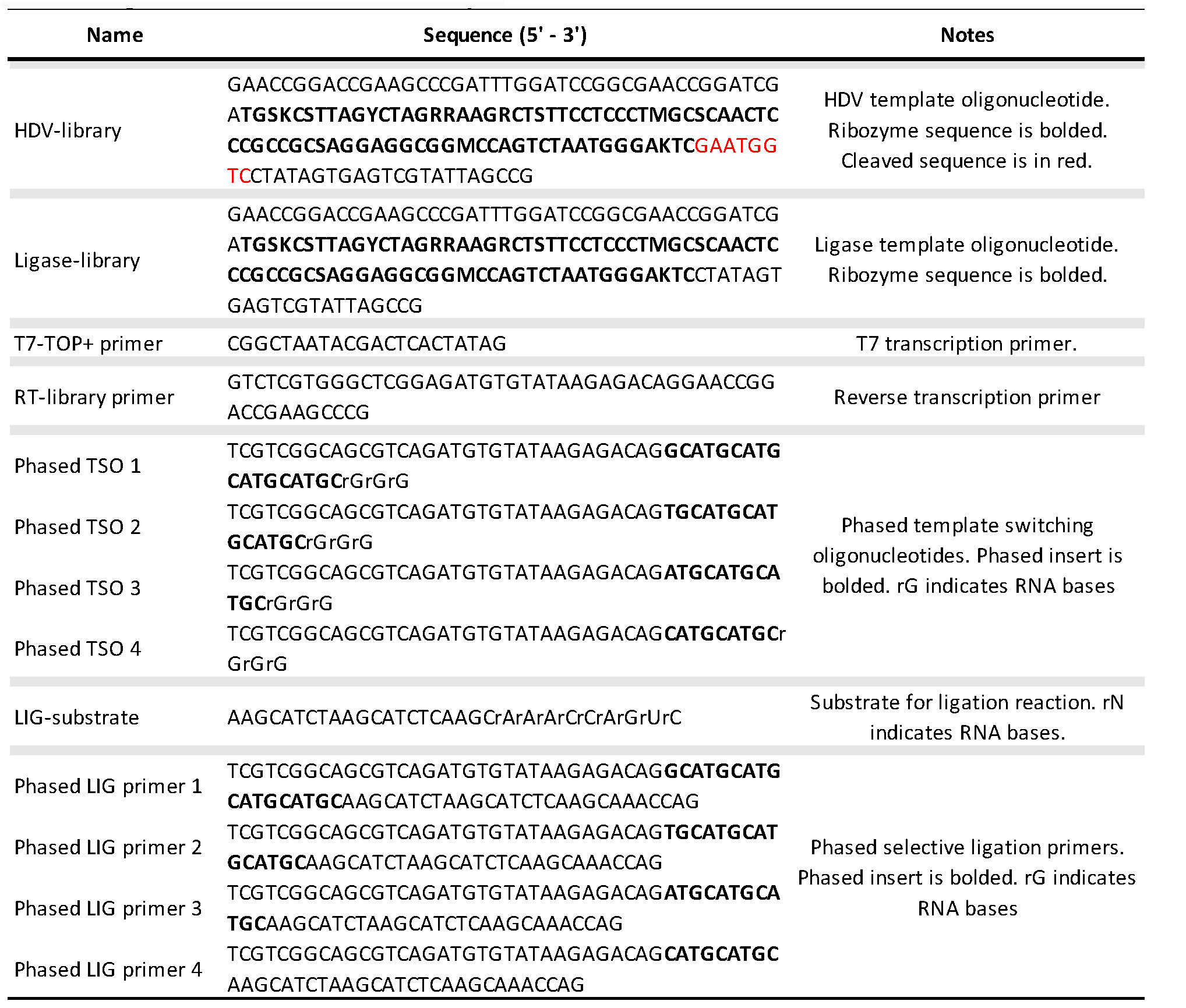
Oligonucleotides used in this study.

### Co-transcriptional Cleavage Assay

The sample preparation was done entirely in triplicate yielding three biological replicates. The ssDNA ultramer cleavage library used for in vitro transcription of the ribozyme mutants was annealed to the T7-TOP+ primer. 20 picomoles each of DNA template and primer were heated for 5 mins at 98°C in 10 μL final volume of custom T7 Mg10 buffer (500 μL 1M Tris pH 7.5, 50 μL 1M DTT, 20 μL 1M Spermidine, 100 μL 1M MgCl_2_, 330 μL RNase-free water). The template and primer were then diluted 10-fold and cooled to room temperature. 2 μL of template and primer were then transcribed in vitro in a 50 μL reaction with 5 μL T7 Mg10 buffer, 1 μL rNTP (25 mM, NEB), 1 μL T7 RNA polymerase (200 units, Thermo Scientific) and 41 μL RNase free water (Ambion) at 37°C for 20 mins. The transcription was then terminated by adding 15 μL of 50 mM EDTA. Although the total amount of cleaved RNA increases during transcription, the ratio of cleaved to uncleaved remains the same, as long as the rate of transcription is constant, which is true for moderately short transcription times before reagents become limited (*39*). 20 mins was determined to be the optimal time for transcription by transcribing the library at multiple time points and measuring RNA levels using denaturing PAGE (Fig. S8A). 20 mins was selected as optimal because it was still during linear growth before reaching a plateau. The transcription reaction was then cleaned and concentrated with Direct-zol RNA MicroPrep w/ TRI-Reagent (Zymo Research) to 7 μL. The concentration of the RNA sample was then determined using a spectrophotometer (ThermoFisher NanoDrop) and the samples were normalized to 5 μM. The transcribed and cleaned RNA (5 picomoles) was mixed with 20 picomoles of RT-library primer (Table 1) in a volume of 10 μL and was heated at 72 °C for 3 mins and then cooled on ice. 4 μL SMARTScribe 5x First-Strand Buffer (Clontech), 2 μL dNTP (10 mM), 2 μL DTT (20 mM), 2 μL phased template switching oligo mix (10 μM), 1 μL water and 1 μL SMARTScribe Reverse Transcriptase (10 units, Clontech) were then added to the RNA template and RT primer. The phased template switching oligo mix consisted of four oligonucleotides that were phased by the addition of 9, 12, 15 or 18 nucleotides (Table 1). The mixture was then incubated at 42 °C for 90 mins. The reaction was stopped and the RNA degraded by heating the sample to 72 °C for 15 mins. The cDNA was then purified using DNA Clean & Concentrator-5 (Zymo Research) and eluted into 7 μL water.

### Ligation Assay

The ssDNA ultramer ligation library used for in vitro transcription of the ribozyme mutants was annealed to the T7-TOP+ primer. 20 picomoles each of DNA template and primer were heated for 5 mins at 98°C in 10 μL water. The template and primer were then transcribed in vitro in a 30 μL reaction with 12 μL rNTP (25mM, NEB), 3 μL MEGAshortscript T7 Reaction Buffer (10X, Thermo Fisher) and 3 μL MEGAshortscript T7 RNA Polymerase (Thermo Fisher) at 37 °C for 2 hours. The DNA was then degraded using 2 μL TURBO DNase (2 units/μL, Thermo Fisher) and incubating at 37 °C for 15 mins. The transcription reaction was then cleaned and concentrated with Direct-zol RNA MicroPrep w/ TRI-Reagent (Zymo Research) to 7 μL. The concentration of the RNA sample was then determined using a spectrophotometer (ThermoFisher NanoDrop) and the samples were normalized to 5 μM. To assess the starting abundance of each genotype prior to in vitro selection, a portion of each sample was aliquoted and reverse transcribed using the template switching protocol identical to what was used for the HDV-library. The transcribed and cleaned RNA (25 picomoles) was mixed with 200mM Tris pH 7.5 in a volume of 10 μL and heated at 65 °C for 2 minutes and then cooled to room temperature. 500 picomoles of ligation substrate (Table 1) were then added with 4 μL MgCl_2_ (50mM) for a total volume of 20 μL. The mixture was then incubated for 2 hours at 37 °C. To reverse transcribe the samples, 10 μL of the ligation reaction were heated with 40 picomoles of RT-library primer and heated to 72 °C for 3 mins and then cooled on ice. 4 μL SMARTScribe 5x First-Strand Buffer (Clontech), 2 μL dNTP (10 mM), 2 μL DTT (20 mM), 1 μL water and 1 μL SMARTScribe Reverse Transcriptase (10 units, Clontech) were then added to the RNA template and RT primer. The mixture was then incubated at 42 °C for 90 mins. The reaction was stopped and the RNA degraded by heating the sample to 72 °C for 15 mins. The cDNA was then purified using DNA Clean & Concentrator-5 (Zymo Research) and eluted into 10 μL water. To amplify the cDNA that had performed the ligation reaction a mix of phased selective ligation PCR primers were used. The PCR reaction consisted of 1 μL purified cDNA, 12.5 μL KAPA HiFi HotStart ReadyMix (2X, KAPA Biosystems), 2.5 μL selective ligation primer, 2.5 μL RT primer and 5 μL water. To prevent bias during the PCR amplification, multiple cycles of PCR were examined using gel electrophoresis and an appropriate PCR cycle was chosen because it was still in linear growth (Fig S8B). Each PCR cycle consisted of 98 °C for 10 s, 63 °C for 30 s and 72 °C for 30 s. The PCR cDNA product was then cleaned using DNA Clean & Concentrator-5 (Zymo Research) and eluted in 12 μL water.

### Illumina adapter PCR

In preparation for high-throughput sequencing, Illumina adapter sequences were added to the cDNA using PCR. Each of the nine samples (3 HDV, 3 ligated, 3 unligated) were each assigned a unique combination of The PCR reaction consisted of 1 μL purified cDNA, 12.5 μL KAPA HiFi HotStart ReadyMix (2X, KAPA Biosystems), 2.5 μL forward, 2.5 μL reverse primer (Illumina Nextera Index Kit) and 5 μL water. To prevent bias during the PCR amplification, multiple cycles of PCR were examined using gel electrophoresis and an appropriate PCR cycle was chosen because it was still in linear growth (Fig. S8B). Each PCR cycle consisted of 98 °C for 10 s, 63 °C for 30 s and 72 °C for 30 s. The PCR cDNA product was then cleaned using DNA Clean & Concentrator-5 (Zymo Research) and eluted in 30 μL water. The final product was then verified using gel electrophoresis.

### High-Throughput Sequencing

In preparation for high-throughput sequencing, the three cleavage replicates, three ligated replicates and three unligated replicates each with unique Illumina adapter barcodes were pooled and sent to the University of Oregon Genomics and Cell Characterization Core Facility. The samples were sequenced using Illumina NextSeq 500 Single End 150 with 25% PhiX addition. This generated ∼125 million reads (Cluster PF Yield) across the nine samples.

### Data Analysis

Sequencing data were analyzed using custom Python scripts. These scripts identified a universally conserved 3’ handle, determined the reacted state (ligated/ unligated or cleaved/ uncleaved) and isolated the 14 mutational nucleotides to determine genotype. This process was repeated for each experimental replicate. A genotype was considered to be a part of the corresponding genotype network only if detected as catalytically active in all three replicates and had a catalytic rate above the uncatalyzed cleavage or ligation rate. The uncatalyzed cleavage rate is estimated to be 7e-7 min^-1^(*40*). The rates of template-directed, nonenzymatic oligonucleotide ligation are estimated to 2.4e-10 min^-1^ for 2’,5’-linkage and 1.5e-8 min^-1^ for 3’,5’-linkage (*41, 42*). Correlation coefficients were determined between pairs of replicates (Fig. S9A). The distribution of HDV and Ligase sequencing read counts were also determined to verify sequencing quality (Fig. S9B).

### Fitness calculations from sequence data

Fitness values for each genotype were determined from the sequence data. Fitness values for the HDV genotypes were calculated from the fraction of each genotype found in the cleaved form divided by the total reads of that genotype in that sample. These fraction cleaved values were normalized by dividing by the fraction cleaved for the HDV reference genotype, resulting in the *HDV fitness* values reported. The Ligase fitness was determined by the level of enrichment following a round of selection for ligase activity. The relative abundance of each genotype was determined by dividing the reads corresponding to that genotype by the total number of reads in that replicate sample. The change in abundance was determined by taking the relative abundance of a specific genotype in the sample selected for ligation activity and dividing it by the relative abundance in the initial library before selection. This value was normalized by dividing by the change in abundance for the Ligase reference sequence, resulting in the *Ligase Fitness* values reported. We observed detectable Ligase activity for all 16,384 sequences. We note that even the lowest fitness Ligase genotypes were still observed as ligated more than 4 separate times in a given replicate, and more than 32 times across all three replicates. We detected HDV activity for 9,032 of the sequences. The least frequent genotypes in our data that showed HDV activity were observed as cleaved more than once in all three replicates, and uncleaved more than 108 times. Genotypes that were not detected as cleaved in a single replicate were not considered active. This approach provides a conservative estimate for genotypes belonging to a given network. Several fitness values from high-throughput sequencing were then compared to values from gel-based assay for fitness validation (fig. S8c).

### Genotype Network and Fitness Landscape Construction

Visualizations of fitness landscapes were constructed using Gephi (*43*). Each node represents a unique genotype and edges connecting genotypes represent a single mutation. ForceAtlas 2 was used to approximate genotype repulsion using a Barnes-Hut calculation. The z-axis in the fitness landscape (Fig. 3A) was generated using the Network Splitter 3D plugin. Peaks in each fitness landscape were defined as genotypes that were surrounded by mutational neighbors with lower relative fitness. This calculation incorporated the measurement error (delta) between replicates. Ruggedness for each landscape was calculated as the average number of peaks within subgraphs (*44*). Each subgraph contains four mutational positions with 16 genotypes and every possible subgraph within each landscape was assessed for peaks. Pairwise epistasis was calculated as *ε =* log10 (*W*_AB_\**W*_wt_ / *W*_A_\**W*_B_), where *W*_A_ and *W*_B_ are the fitness of RNA variants with a single mutation, *W*_AB_ is the fitness of the variant with both mutations, and *W*_wt_ is the fitness of the wild-type (*6*). Epistasis was calculated for every subgraph of two mutational positions containing four genotypes.

### Evolutionary Simulations

Computational simulations of evolution were accomplished using custom Python scripts that model evolution based on the Wright-Fisher approach (*26, 45*). Simulation started with 1000 individuals of the same genotype. Every generation (update) a new population of 1000 genotypes was generated in the following way. First, a parent genotype from the population was selected at random. The fitness of the genotype was compared to a randomly selected value from a fitness range (between 0 and 1). If the genotype fitness was less than the random value, the genotype was not placed in the new generation. If the genotype fitness was greater than or equal to the random value, it was placed in the new generation, with a chance of mutating at a single, randomly chosen nucleotide position. Mutations occurred if a randomly generated number was lower than the mutation rate set at the beginning of the simulation and remained constant (μ = 0.01). This process was repeated until 1000 individuals were placed in the new generation. The simulation then repeated this process for 1000 generations. We carried out the simulations on the Ligase landscape starting from the 17 genotypes with HDV fitness ≥ 1 and did so for a total of 100 replicates for each genotype (Fig. S2). The 100 replicates for each starting genotype were averaged (Fig 4b) and the initial rate of adaptation and unique genotypes explored for each starting genotype were calculated. For each simulation, *initial rate* was determined by subtracting the population fitness at *generation* = 0 from the population fitness at *generation* = 200 and dividing this value by 200 *generations*. We also ran simulations on the HDV landscape starting from the 17 genotypes with the highest Ligase genotypes that also had non-zero HDV fitness. These were repeated for 100 replicates (Fig. S5) and were averaged (Fig. 4E). Initial rate was also calculated for these simulations (Fig. 4E). Lastly, simulations were conducted on a HDV-Ligase co-selection landscape that allows selection to act upon both functions simultaneously. For this model, the fitness was calculated as: *W*_*HDV*_\**β*_*HDV*_+ *W*_*Ligase*_\**β*_*Ligase*_, where *W* indicates the fitness of that function and *β* indicates a weighting parameter that can be adjusted (Fig. S6–S7). Otherwise, simulations were the same as above.

**Fig. 4.**
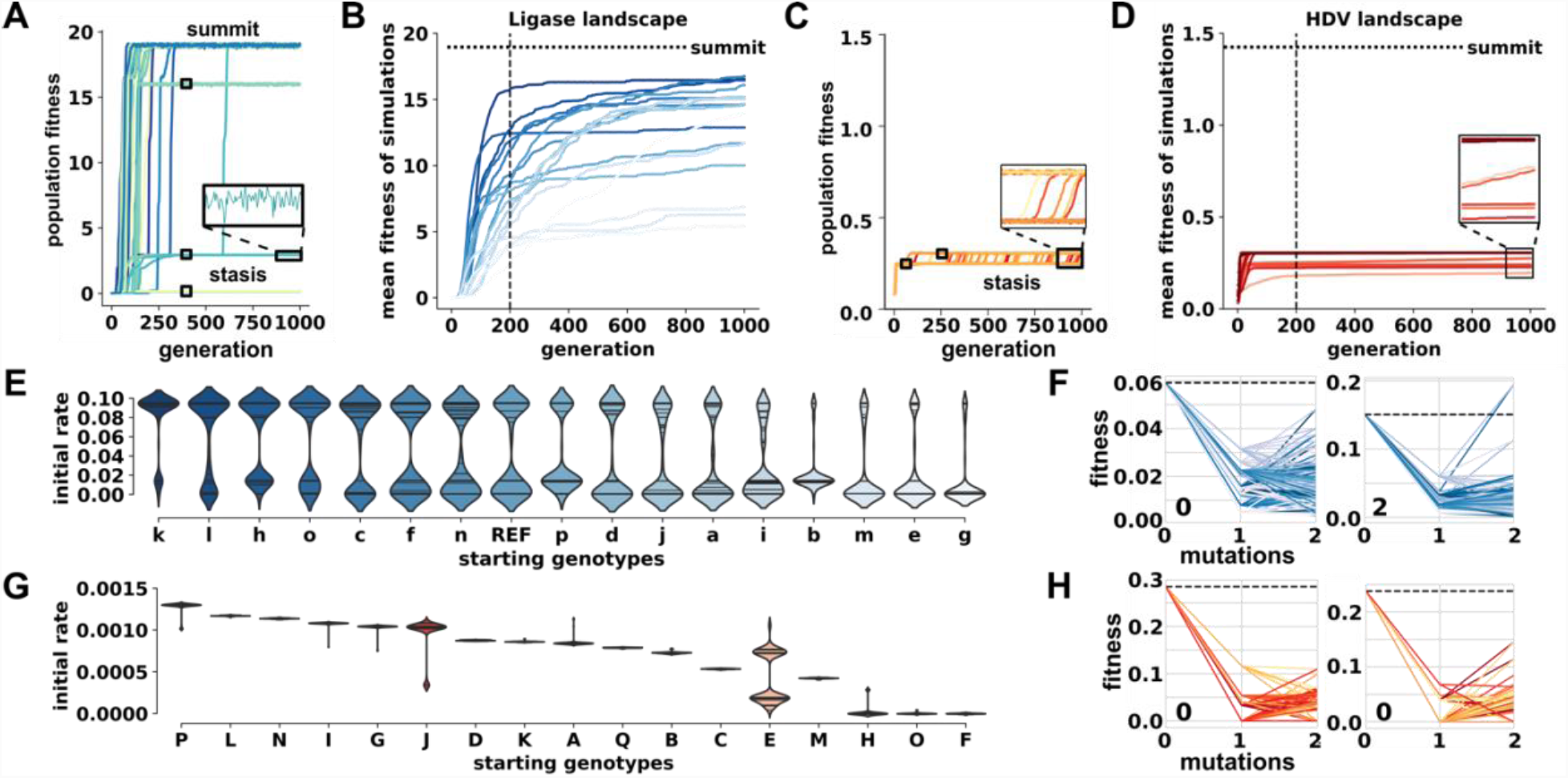
Starting genotypes result in different rates of evolutionary adaptation. **(*A*)** Rates of Ligase adaptation from a single HDV genotype. Each trace shows the average *population fitness* as a function of generation time for a separate simulation of 1000 individuals each. The traces from 100 separate simulations are shown. Inset shows minor fluctuations during periods of stasis. **(*B*)** Average rates of evolutionary adaptation of Ligase activity starting from 17 genotypes. Each trace represents a different starting genotype (*a-p* and *HDV reference*) and shows the mean fitness of 100 simulations as a function of time (*generation*). The y-axis is scaled to the maximum fitness on this landscape (*summit*, horizontal dashed line). The vertical dashed line marks generation 200. **(*C*)** Rates of HDV adaptation from a single Ligase genotype. **(*D*)** Average rates of evolutionary adaptation of HDV activity starting from 17 genotypes. Each trace represents a different starting genotype (*A-Q)* and shows the mean fitness of 100 simulations as a function of time (*generation*). The y-axis is scaled to the maximum fitness on this landscape (*summit*, horizontal dashed line) **(*E*)** Distributions of initial rates of adaptation during simulations on the Ligase landscape. Initial rate is determined as the population fitness divided by the generations at 200 generations. Each violin plot represents the distribution of 100 simulations starting from the same genotype, which is indicated on the x-axis. **(*F*)** Sign epistasis in the local fitness landscape of genotypes that cause periods of stasis in the Ligase landscape. The fitness of the stasis genotype is plotted at *mutations* = 0 and this starting fitness is marked with a dashed line. The fitness of neighboring genotypes that differ by 1 or 2 mutations are shown. Distributions of initial rates of adaptation during simulations on the HDV landscape. **(*G*)** Distributions of initial rates of adaptation during simulations on the HDV landscape. **(*H*)** Sign epistasis in the local fitness landscape of genotypes that cause periods of stasis in the HDV landscape.

## Supporting information

## General

We would like to thank A. Wagner (University of Zurich) and J. Payne (ETH) for helpful discussions.

## Author contributions

Conceptualization – EJH, Methodology – DPB JC, Software – DPB BØ, Formal Analysis – EJH DPB BØ, Investigation – DPB JC, Writing (Original Draft Preparation) – EJH DPB, (Review and Editing) – EJH DPB JC BØ, Visualization – DPB EJH.

## Data and materials availability

Genotype and fitness values are available as SI Dataset. Raw sequence data, code and materials are available upon request.

## Supplementary Materials

Fig. S1 Overlay of HDV and Ligase genotype networks with varying fitness cut-offs.

Fig. S2 Rate of adaptation for populations starting from different genotypes on the Ligase landscape.

Fig. S3 Trajectories away from *stasis genotypes*.

Fig. S4 Characterization of *stasis genotype* I.

Fig. S5 Rate of adaptation for populations starting from different genotypes on the HDV landscape.

Fig. S6 Evolutionary simulations on the HDV-Ligase co-select fitness landscapes.

Fig. S7 Individual traces of evolutionary adaptation on HDV-Ligase co-select fitness landscapes.

Fig. S8 Time-courses for sample optimization and validation of sequencing fitness values.

Fig. S9 High-throughput sequencing results for HDV and Ligase.

Table S1. Starting Genotypes used in Evolution Simulations.

Data file S1 HDV-Ligase fitness measurements from high-throughput sequencing assays.

Movie S1 Aerial overview of the HDV-Ligase fitness landscape.

Movie S2 Simulated evolution on Ligase landscape starting at genotype k.

Movie S3 Simulated evolution on Ligase landscape starting at genotype a.

Movie S4 Simulated evolution on Ligase landscape starting at genotype b.

Movie S5 Simulated evolution on Ligase landscape starting at genotype m.

Movie S6 Simulated evolution on HDV landscape starting at genotype A.

**Data file S1 HDV-Ligase fitness measurements from high-throughput sequencing assays.** The fitness measurements for each of the 16,384 unique genotypes presented in this study. The genotypes are displayed as the 14 mutational positions. HDV and Ligase fitness values are colored according to the relative fitness for easier interpretation. Delta values were calculated as the standard error between the three sequencing replicates for each function.

**Movie S1 Aerial overview of the HDV-Ligase fitness landscape.** Overview of the empirical HDV-Ligase fitness landscape presented in Fig. 3A. Each node represents an individual genotype and edges connect nodes that differ by a single nucleotide. The size and height of each node indicates the relative genotype fitness (HDV-red, Ligase-blue).

**Movie S2 Simulated evolution on Ligase landscape starting at genotype k.** White nodes are genotypes in the fitness landscape. Genotypes are connected by light blue edges if they differ by a single nucleotide change. The size of the blue circles depicts the relative proportion of the simulated population at that genotype. The y-axis is relative Ligase fitness. The x-axis is number of nucleotide differences from the HDV reference sequence (mutational distance). The number above the graph represents the generation number. The population average is also depicted with the number of generations on the x-axis and mean population fitness on the y-axis. Lastly, the population diversity at a given generation is plotted as a function of generational time. Population diversity indicates the number of unique genotypes present in the population.

**Movie S3 Simulated evolution on Ligase landscape starting at genotype a.** White nodes are genotypes in the fitness landscape. Genotypes are connected by light blue edges if they differ by a single nucleotide change. The size of the blue circles depicts the relative proportion of the simulated population at that genotype. The y-axis is relative Ligase fitness. The x-axis is number of nucleotide differences from the HDV reference sequence (mutational distance). The number above the graph represents the generation number. The population average is also depicted with the number of generations on the x-axis and mean population fitness on the y-axis. Lastly, the population diversity at a given generation is plotted as a function of generational time. Population diversity indicates the number of unique genotypes present in the population.

**Movie S4 Simulated evolution on Ligase landscape starting at genotype b.** White nodes are genotypes in the fitness landscape. Genotypes are connected by light blue edges if they differ by a single nucleotide change. The size of the blue circles depicts the relative proportion of the simulated population at that genotype. The y-axis is relative Ligase fitness. The x-axis is number of nucleotide differences from the HDV reference sequence (mutational distance). The number above the graph represents the generation number. The population average is also depicted with the number of generations on the x-axis and mean population fitness on the y-axis. Lastly, the population diversity at a given generation is plotted as a function of generational time. Population diversity indicates the number of unique genotypes present in the population.

**Movie S5 Simulated evolution on Ligase landscape starting at genotype m.** White nodes are genotypes in the fitness landscape. Genotypes are connected by light blue edges if they differ by a single nucleotide change. The size of the blue circles depicts the relative proportion of the simulated population at that genotype. The y-axis is relative Ligase fitness. The x-axis is number of nucleotide differences from the HDV reference sequence (mutational distance). The number above the graph represents the generation number. The population average is also depicted with the number of generations on the x-axis and mean population fitness on the y-axis. Lastly, the population diversity at a given generation is plotted as a function of generational time. Population diversity indicates the number of unique genotypes present in the population.

**Movie S6 Simulated evolution on HDV landscape starting at genotype A.** White nodes are genotypes in the fitness landscape. Genotypes are connected by light red edges if they differ by a single nucleotide change. The size of the red circles depicts the relative proportion of the simulated population at that genotype. The y-axis is relative HDV fitness. The x-axis is number of nucleotide differences from the HDV reference sequence (mutational distance). The number above the graph represents the generation number. The population average is also depicted with the number of generations on the x-axis and mean population fitness on the y-axis. Lastly, the population diversity at a given generation is plotted as a function of generational time. Population diversity indicates the number of unique genotypes present in the population.

## Supplementary Materials

**Table S1.**
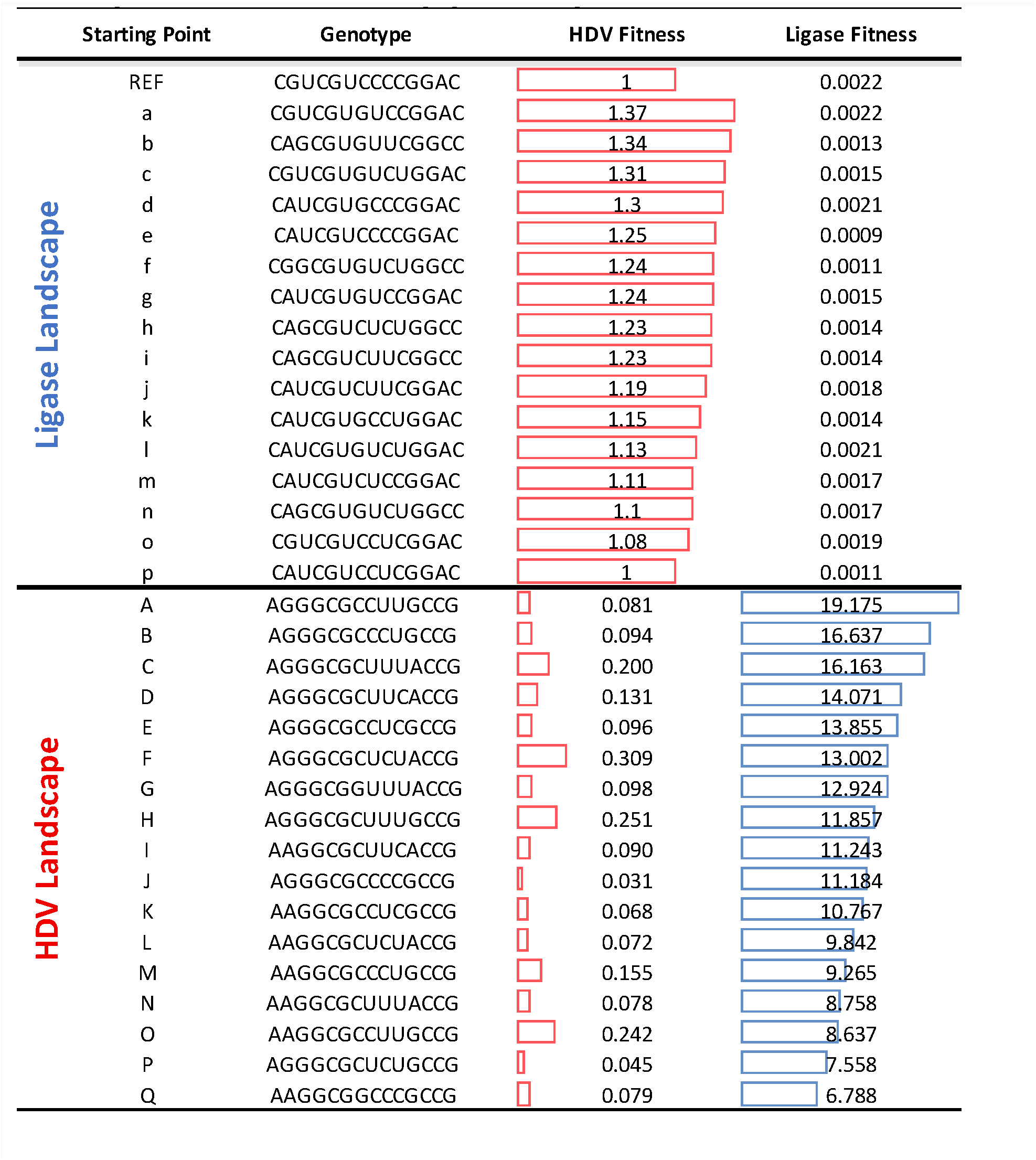
Starting Genotypes used in Evolution Simulations. Genotypes are represented by the unique combination of nucleotides in the 14 variable positions of the library. Starting point letters correspond to Fig. 3A. HDV and Ligase fitness are colored with bar graphs indicating the relative fitness.

**Fig. S1.**
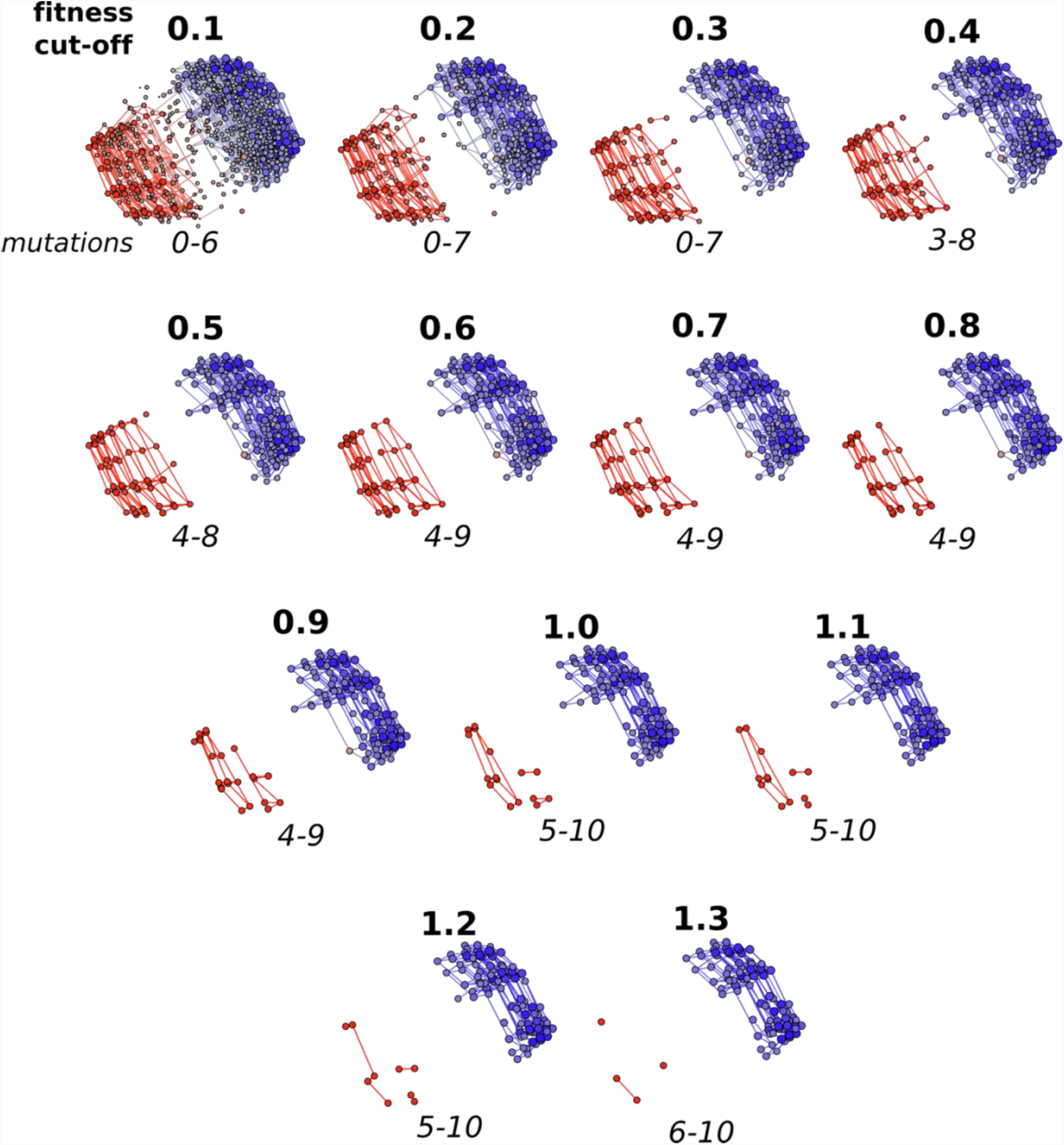
Overlay of HDV and Ligase genotype networks with varying fitness cut-offs. Each plot indicates the overlay of the two networks with all genotypes with fitness values below the cut-off removed. The size of the node indicates relative fitness and nodes are colored based on their dominant activity (red = HDV, blue = Ligase). Below each plot is the range of mutations needed to go from one network to the other. The distribution of the number of mutations required is displayed for each fitness cut-off in Fig. 2A.

**Fig. S2.**
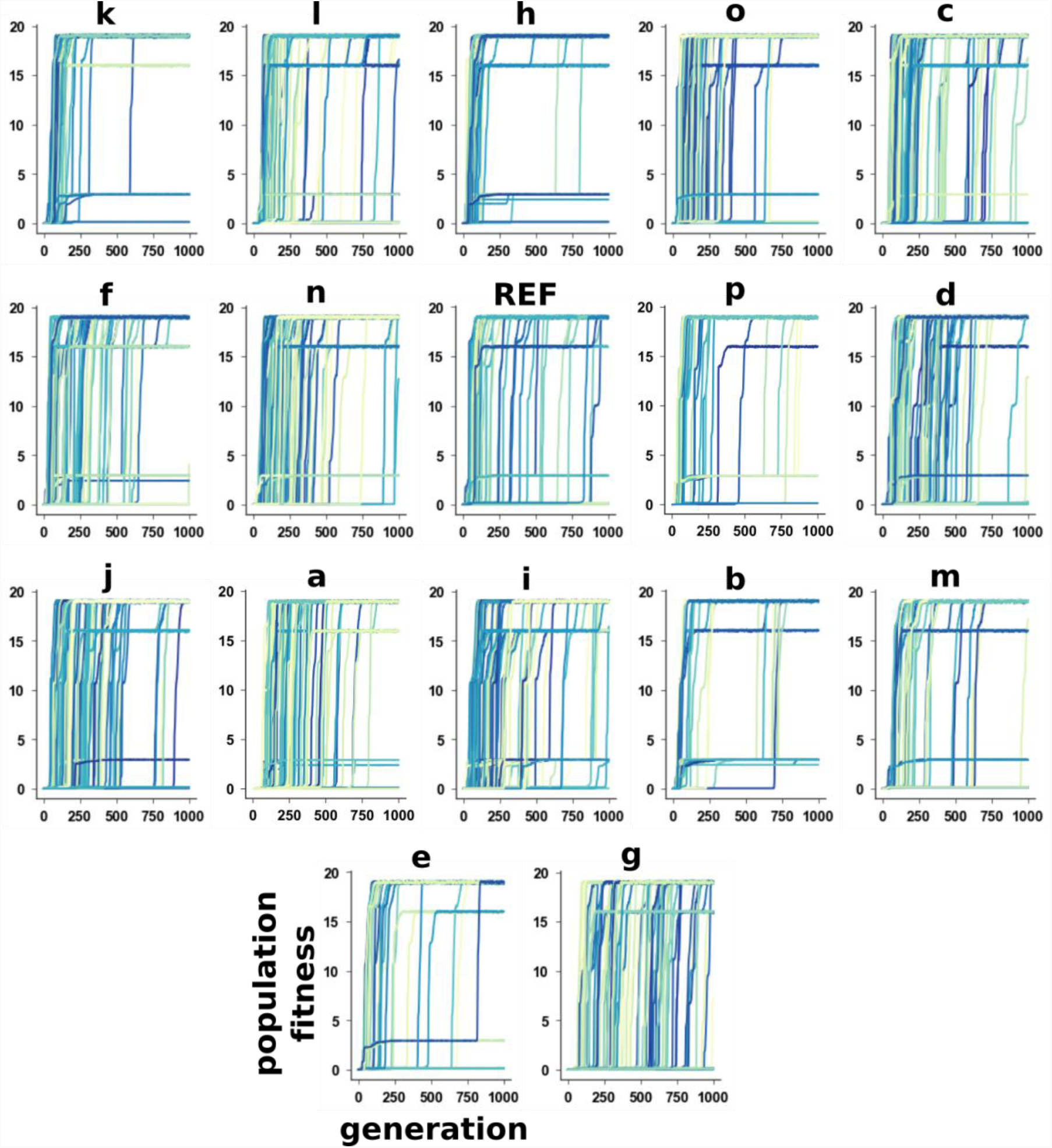
Rate of adaptation for populations starting from different genotypes on the Ligase landscape. Each trace shows the increase in population fitness over generation time for a single simulation of 1000 individuals. Each plot shows 100 simulations starting from the same genotype. All starting genotypes has *HDV fitness* ≥ 1. The letter above each subplot indicates the starting point from the network as shown in Fig. 3a. Letters were assigned alphabetically based on highest to lowest HDV fitness and genotype **a** represents the genotype with the highest measured HDV fitness. The graphs are ordered from fastest to slowest initial rates (Fig. 4D).

**Fig. S3.**
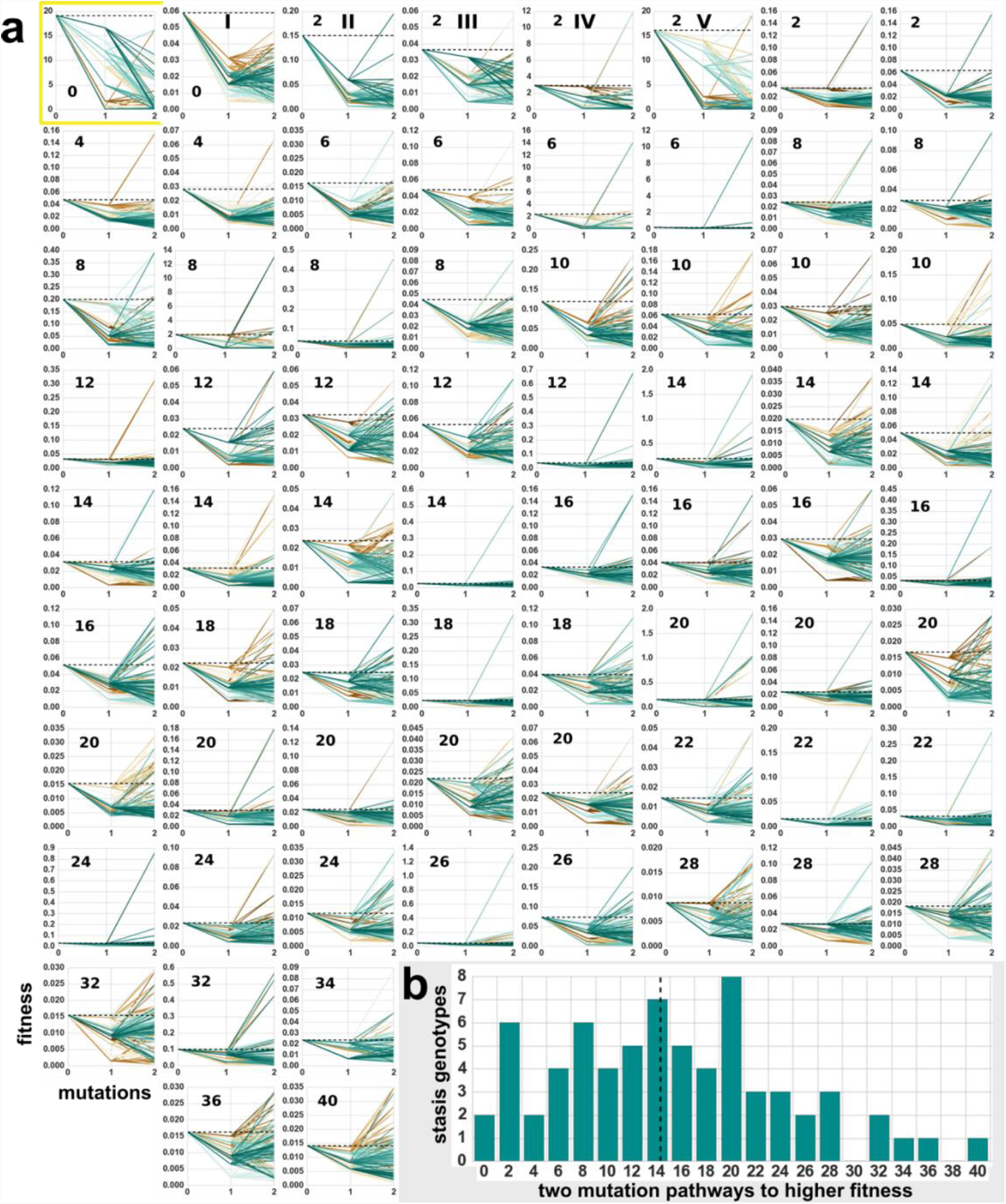
Trajectories away from *stasis genotypes.* **(A)** Each line leads from the stasis genotype (*Mutations* = 0) to one and two mutations away. All 69 stasis genotypes (peaks) in the Ligase fitness landscape are depicted. The number on each graph represents the number of two mutation pathways to higher fitness from each stasis genotype. Yellow box indicates the genotype with the highest measured Ligase fitness. **(B)** The distribution of two mutation pathways to higher fitness genotypes from each stasis genotypes in the Ligase landscape. The dotted vertical line indicates the mean of the distribution.

**Fig. S4.**
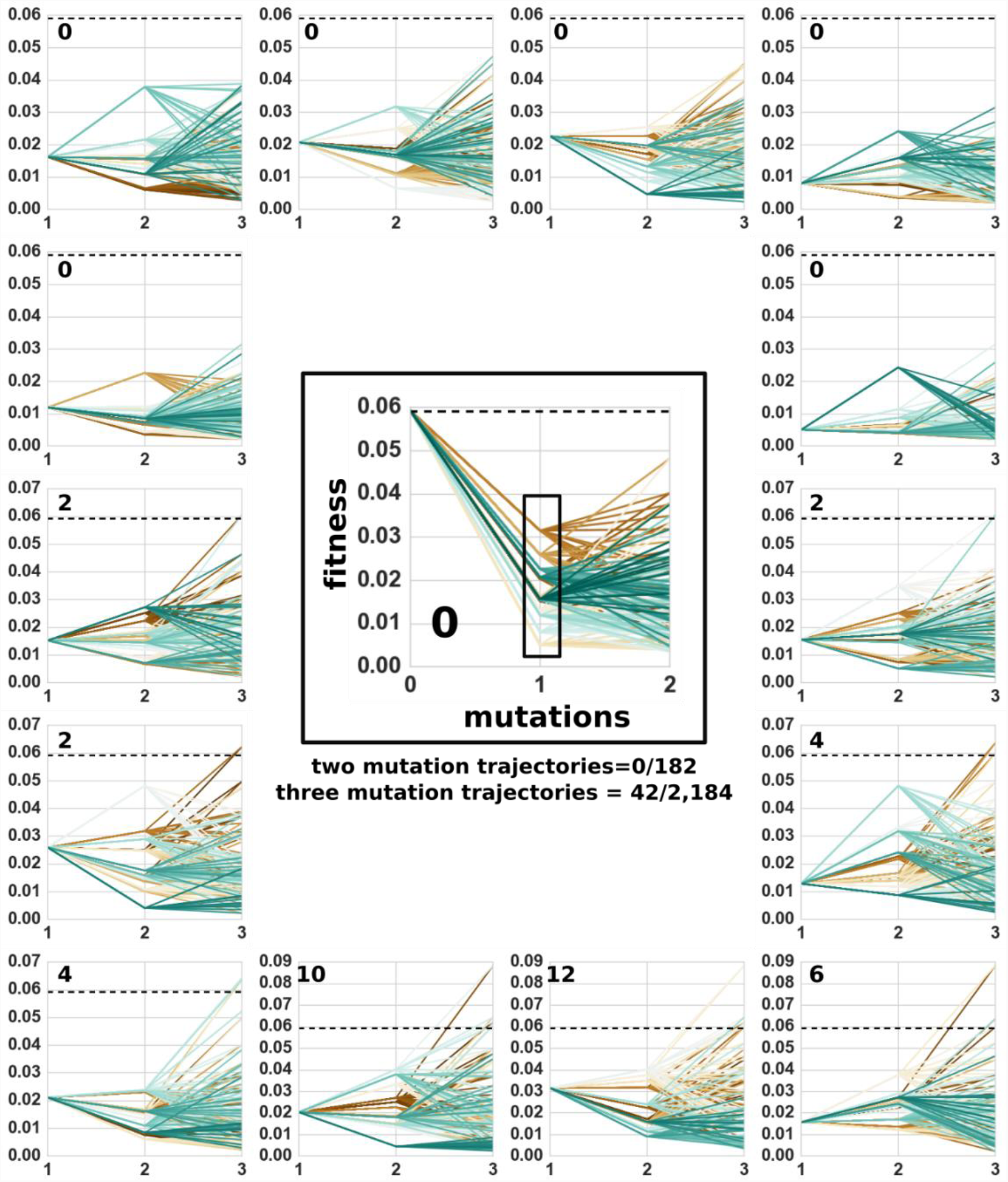
Characterization of *stasis genotype* I. Stasis genotype I from Fig. 4d is depicted in the center with each of the two mutation trajectories. None of the 182 two mutation trajectories lead to higher fitness than the stasis genotype (mutation = 0). The pathways two mutations from each of the 14 genotypes that are a single mutation away from the stasis genotype are individually depicted. In total, 42 out of a possible 2,184 three mutation trajectories yield a higher fitness than the initial stasis genotype (dashed line).

**Fig. S5.**
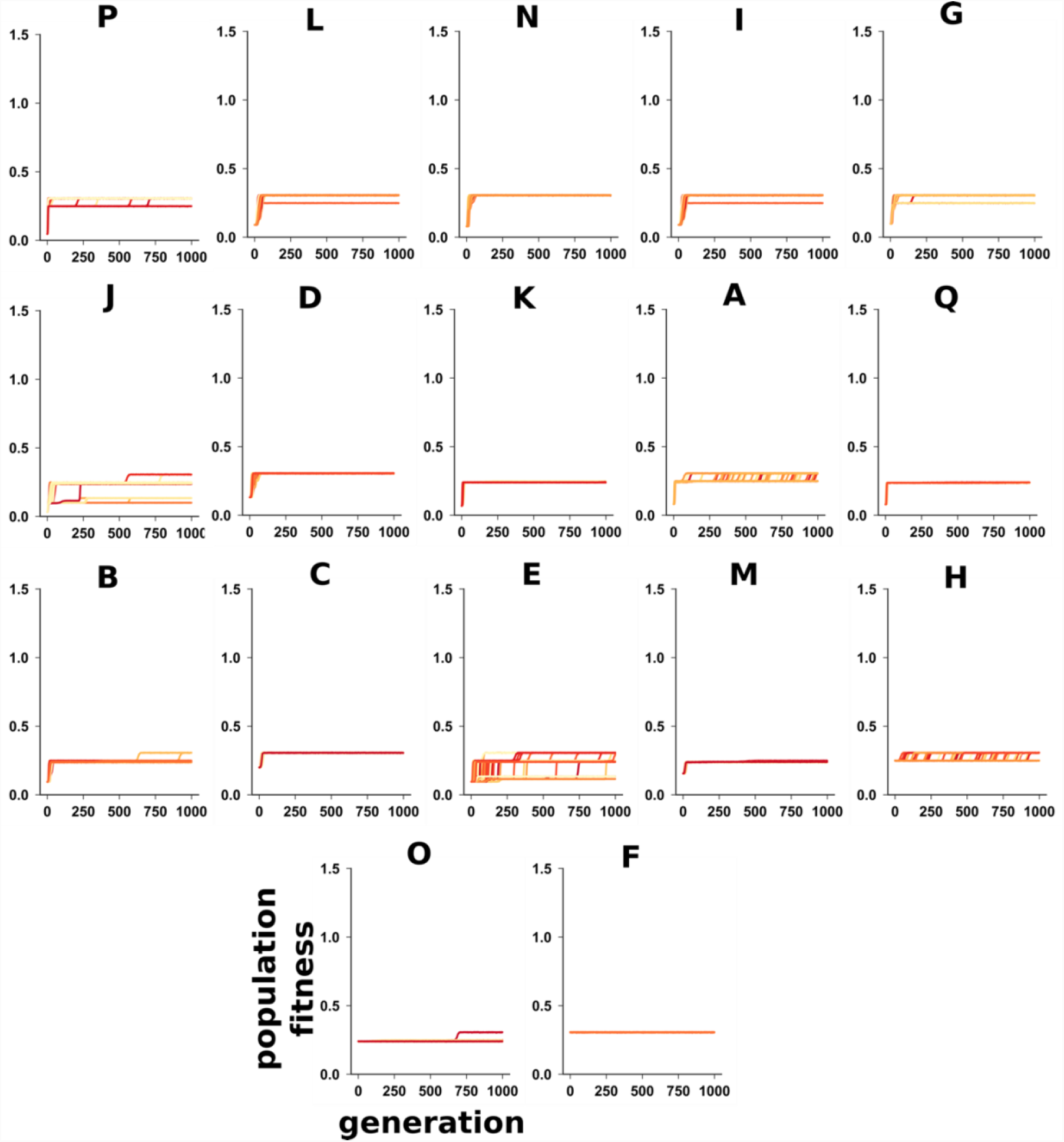
Rate of adaptation for populations starting from different genotypes on the HDV landscape. Each trace shows the increase in population fitness over generation time for a single simulation of 1000 individuals. Each plot shows 100 simulations starting from the same genotype. The letter above each subplot indicates the starting point from the network as shown in Fig. 3a. Letters were assigned alphabetically based on highest to lowest Ligase fitness and genotype **A** represents the genotype with the highest measured Ligase fitness. The graphs are ordered from fastest to slowest initial rates (Fig. 4E).

**Fig. S6.**
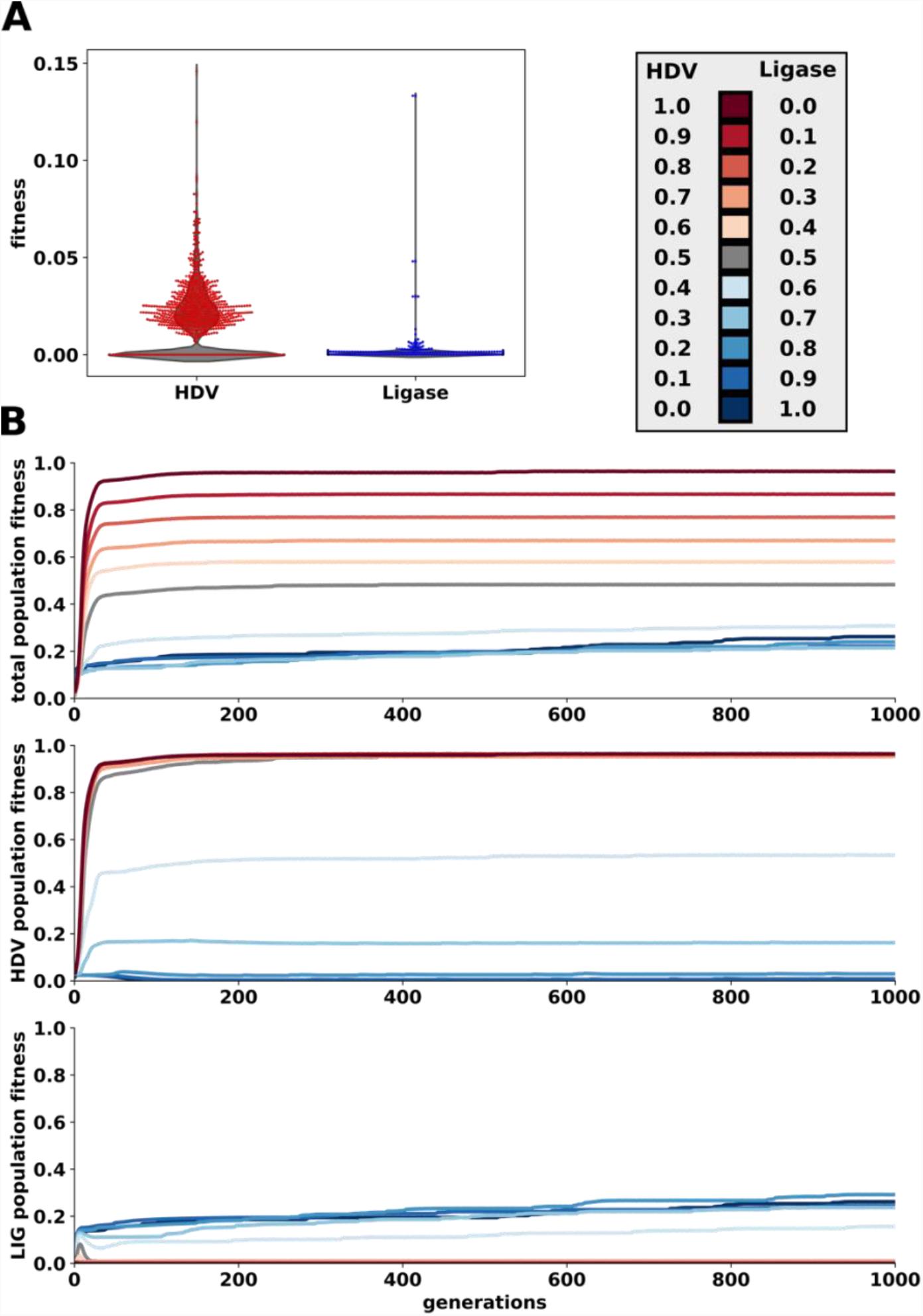
Evolutionary simulations on the HDV-Ligase co-select fitness landscapes. **(A)** The starting population that was randomly selected from the 3,432 genotypes that are 7 mutations from HDV reference and Ligase reference. **(B)** Average rates of evolutionary adaptation on the HDV-Ligase co-select fitness landscapes with varying weighted parameters (β) for each function. Line indicates the average of 100 replicates (Fig. S7). Total population fitness indicates the fitness resulting from the following equation, *W_HDV_*β_HDV_+ W_Ligase_*β_Ligase_*, where *W* indicates the fitness of that function and *β* indicates a weighting parameter that was adjusted. The HDV and Ligase population is also plotted independently to indicate which function is the dominant contributor to the total fitness. Line color indicates the weighting parameters used in the simulation as indicated in the top-right inset.

**Fig. S7.**
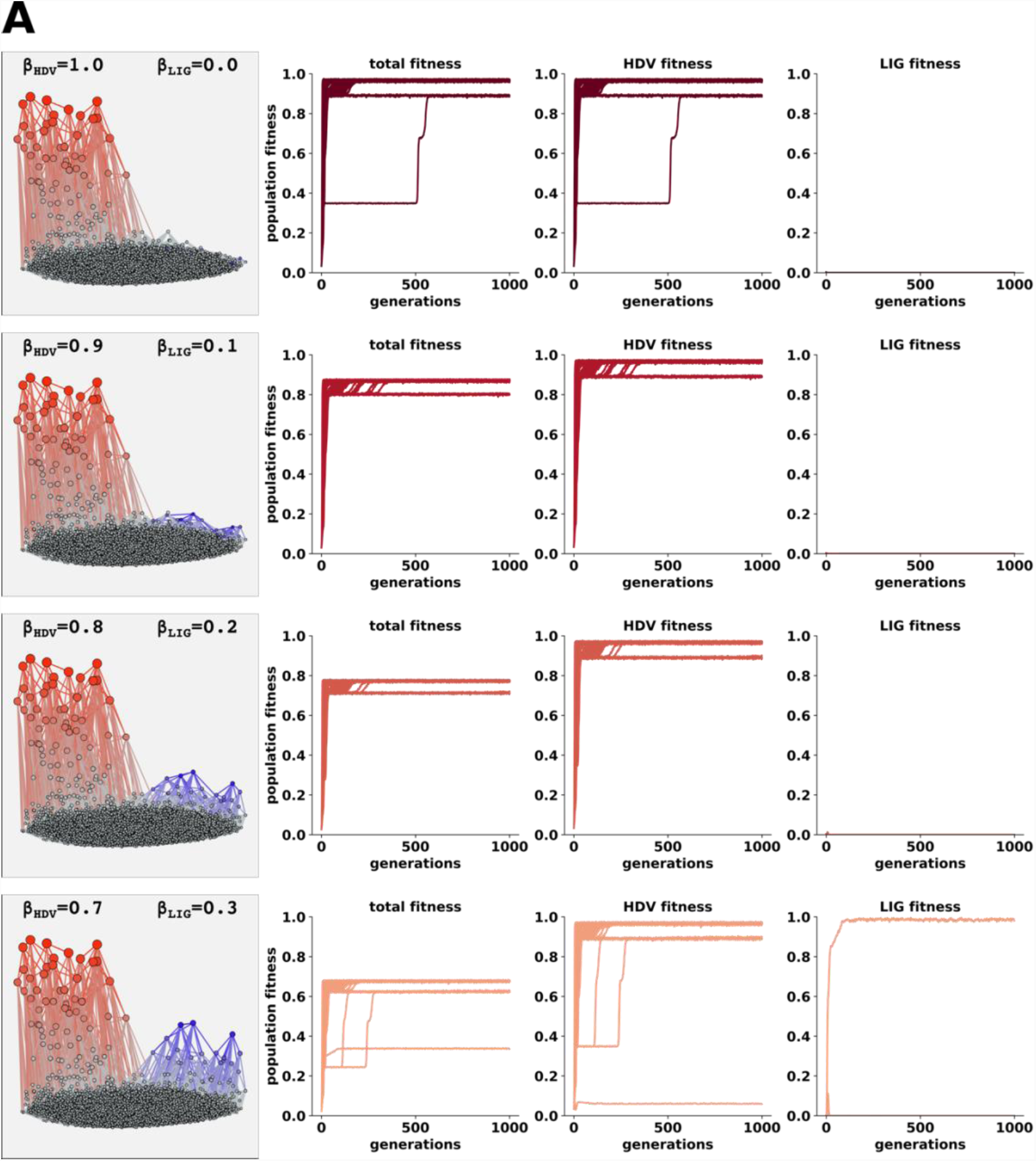
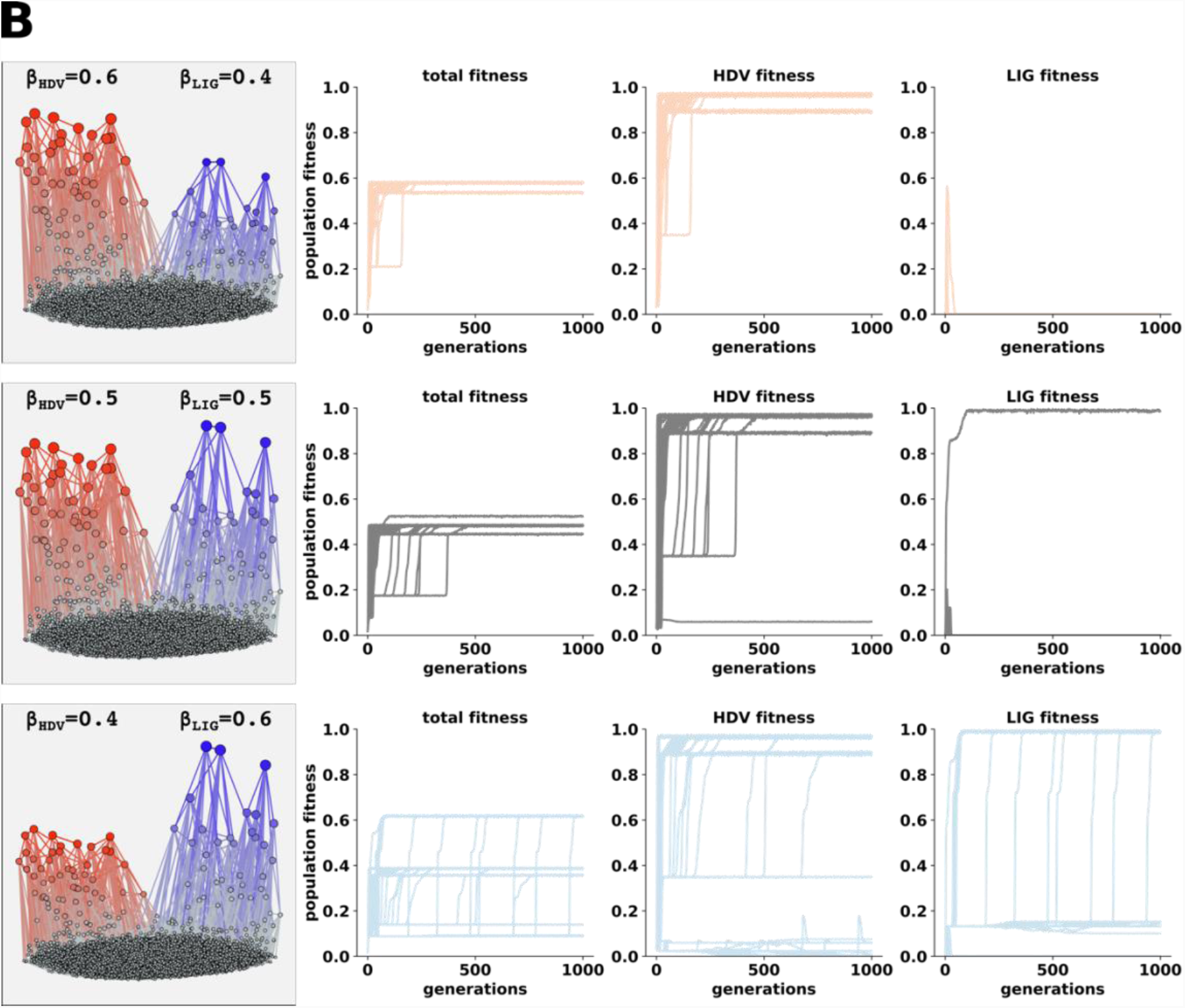
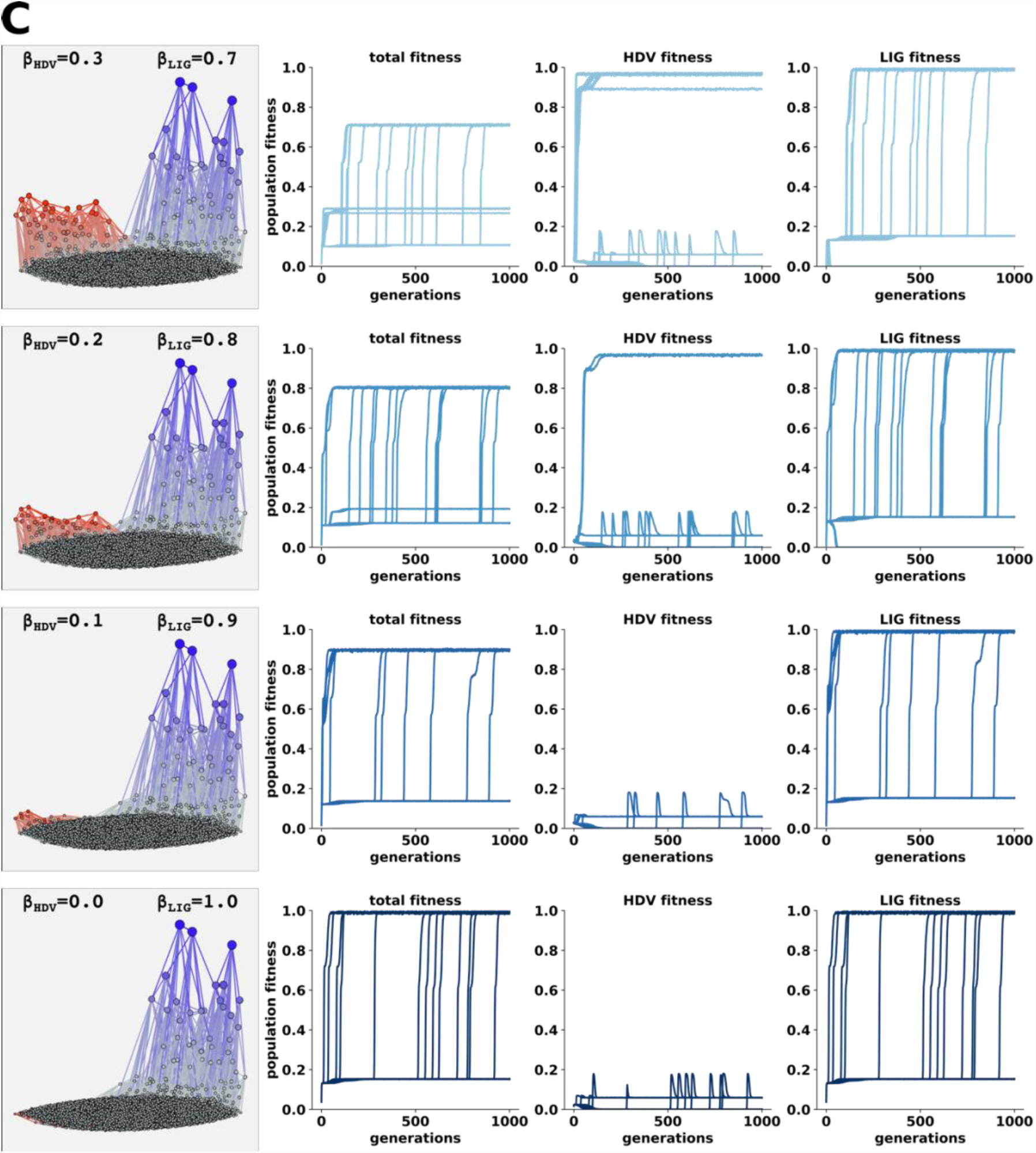
Individual traces of evolutionary adaptation on HDV-Ligase co-select fitness landscapes. **(A-C)** Left plot indicates the architecture of the fitness landscape for each combination of weighting parameters (β). Total fitness (calculated as *WHDV*βHDV+ WLigase*βLigase*), HDV fitness and Ligase fitness are shown for each individual simulation replicate. Color of lines correspond to the weighting parameters discussed in Fig. S6.

**Fig. S8.**
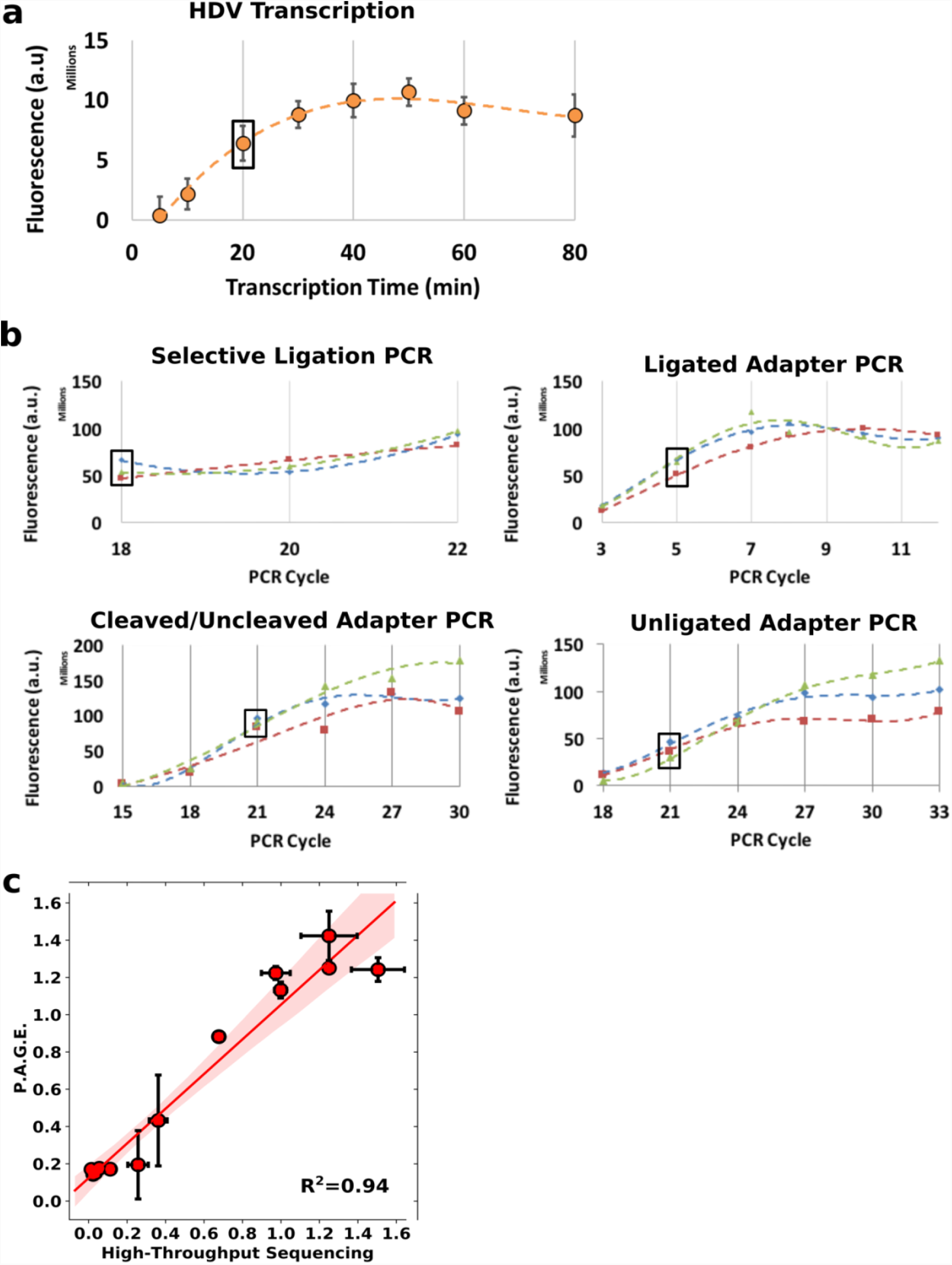
Time-courses for sample optimization and validation of sequencing fitness values. **(A)** Time-course transcription for total RNA yield using the developed co-transcriptional cleavage assay. Data points indicate the mean RNA yield of five replicates. Error bars are standard error of the mean. Samples were run on 10% denaturing polyacrylamide gel, visualized with GelRed (Biotium), and quantified by densitometry. The time chosen as optimal (20 mins) is indicated with a box. **(B)** Time-course PCR was performed for the selective ligation PCR and each Illumina adapter PCR for each replicate (blue, green, red). Samples were run on 2% agarose gel, visualized with GelRed (Biotium), and quantified by densitometry. The black box indicates the PCR cycle that was determined to be optimal for each PCR reaction. **(C)** Correlation of fitness values for thirteen unique ribozyme genotypes assessed by high-throughput sequencing and gel-based assay (P.A.G.E). Both methods assessed the fraction cleaved (fitness) of each genotype.

**Fig. S9.**
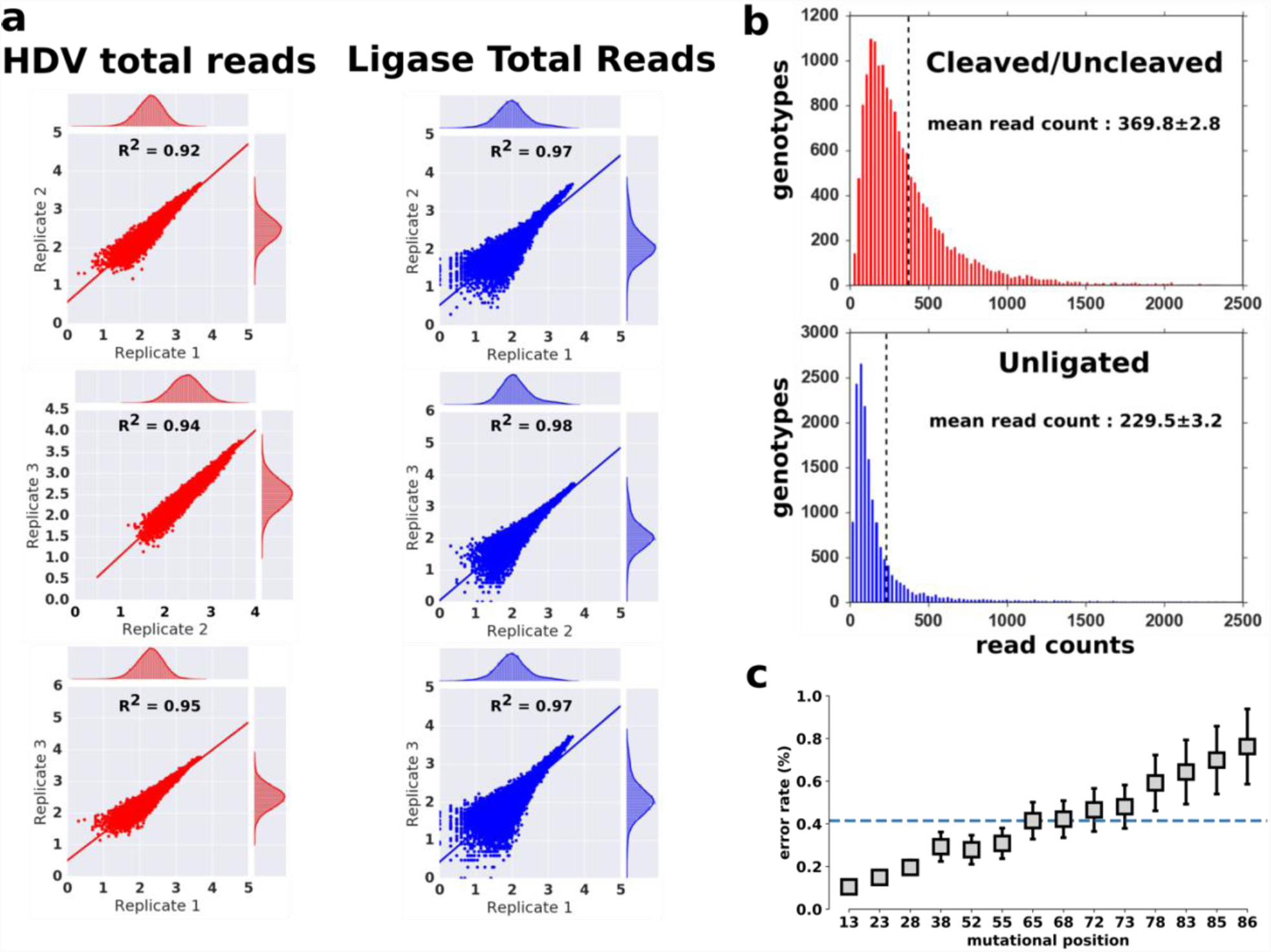
High-throughput sequencing results for HDV and Ligase. **(A)** Correlation of total HDV and Ligase reads for each of the three replicates. Each figure consists of all 16,384 genotypes presented in this study. Each data point represents the frequency that a specific sequence was observed in a particular replicate (y-axis) vs. another replicate (x-axis). Sequence kernel density estimation is also reported from each replicate in the jointplot (Seaborn). The number of reads on the x and y-axis are log10 transformed. **(B)** Distribution of sequencing read counts. Histograms indicating the average read counts for each individual genotype for the HDV and Ligase samples. The mean read count for each genotype in HDV and Ligase replicates was 369 and 230, respectively. **(C)** Error rates calculated from base miscalls in the PhiX reference genome. Error rate (y-axis) is shown for the 14 positions (x-axis) where our genotypes are defined. Each position is read in four different sequencing cycles, and error rates are reported as the average error rate of these four cycles. Dashed blue line indicates the average error rate across all 14 mutational positions. Error rates are calculated by aligning each PhiX sequence read in our data to the reference PhiX genome and counting mismatches at each sequence cycle.

